## Supplemental Data for "Seasonal and comparative evidence of adaptive gene expression in mammalian brain size plasticity"

### **SUPPLEMENTAL FIGURES**

**Supplemental Figure 1.**

(A) Phylogeny used for comparative transcriptomics of the hypothalamus with (B) each species normalized count distribution of single copy orthologs (6496 genes). Visualization shows the efficacy of normalization as each species has roughly similar distributions.

**Supplemental Figure 2.**

Hierarchical clustering of 786 gene expression profiles formed 12 distinct clusters in the shrew hypothalamus. Of these 12 clusters, five clusters (Clusters 2, 3, 8, 11, 12) consisted of 392 genes which resembled a large expression divergence between summer juveniles and the remaining individuals. These genes show expression divergence between recently postnatal shrews and the other stages, compared to the variation found across Dehnel’s phenomenon (Clusters 1, 4, 5, 6, 7, 9, 10).
