## Supplementary figures and images for "Seasonal and comparative evidence of adaptive gene expression in mammalian brain size plasticity"

### Supplemental Figure 1

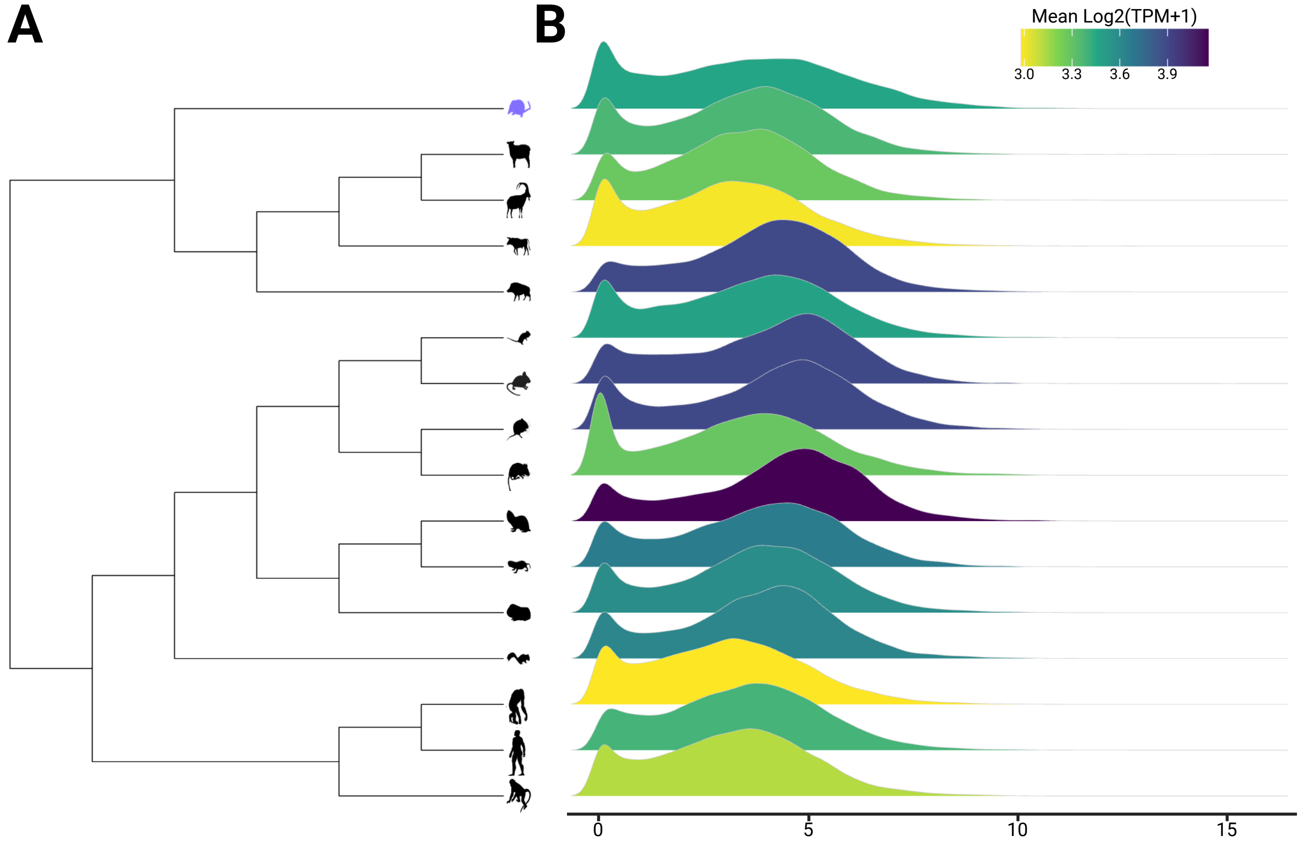

### Supplemental Figure 2

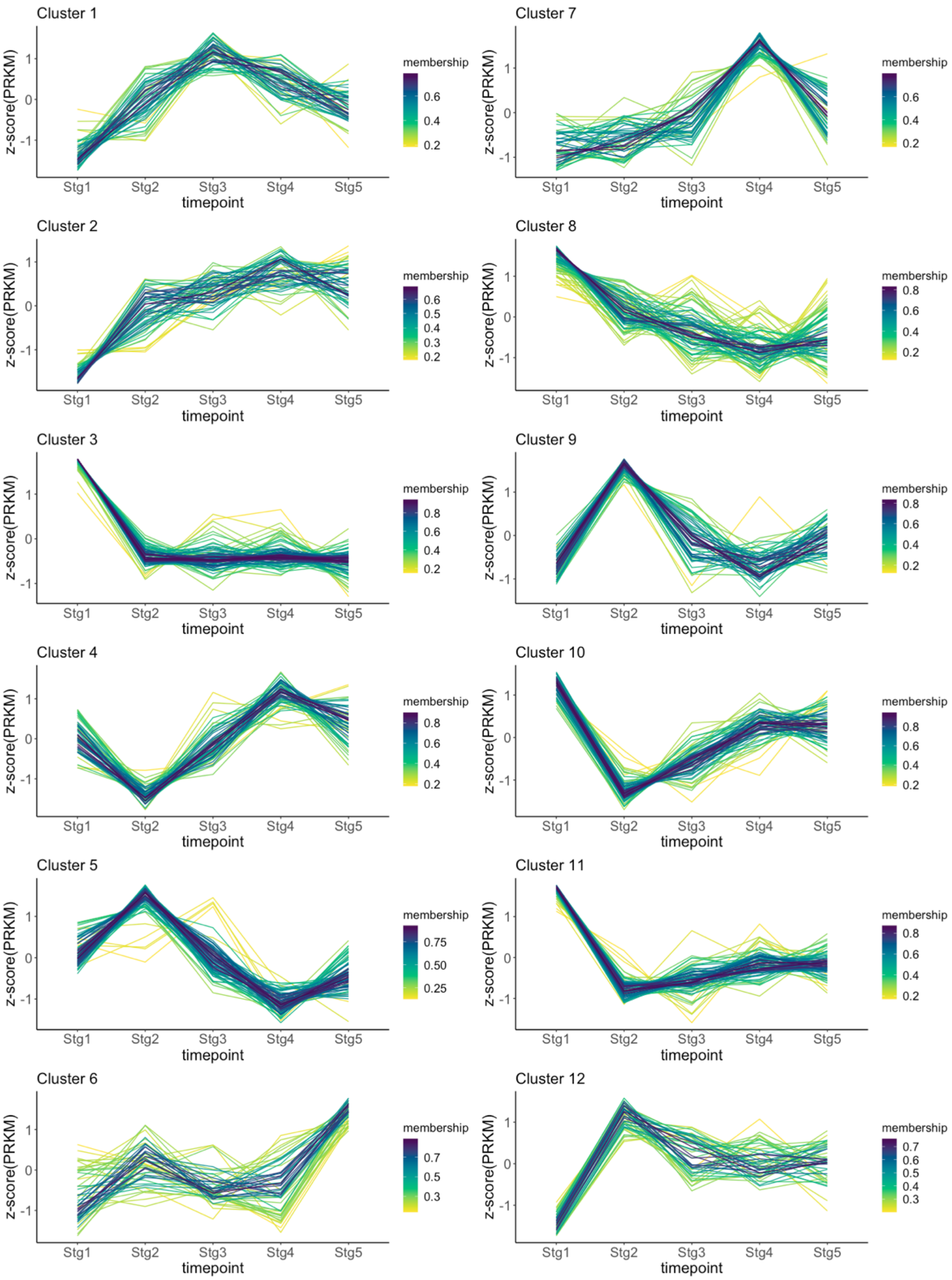
